# NAD^+^ Redox Imbalance in the Heart Exacerbates Diabetic Cardiomyopathy

**DOI:** 10.1101/2020.07.03.183111

**Authors:** Ying Ann Chiao, Akash Deep Chakraborty, Christine M. Light, Rong Tian, Junichi Sadoshima, Xiaojian Shi, Haiwei Gu, Chi Fung Lee

**Affiliations:** Aging and Metabolism Research Program, Oklahoma Medical Research Foundation, Oklahoma City, OK; Cardiovascular Biology Research Program, Oklahoma Medical Research Foundation, Oklahoma City, OK; Mitochondria and Metabolism Center, University of Washington, Seattle, WA; Department of Cell Biology and Molecular Medicine, Rutgers New Jersey Medical School, Newark, NJ; Arizona Metabolomics Laboratory, College of Health Solutions, Arizona State University, Scottsdale, AZ; Department of Physiology, University of Oklahoma Health Sciences Center, Oklahoma City, OK

## Abstract

**Background:** Diabetes is a risk factor of heart failure and promotes cardiac dysfunction. Diabetic tissues are associated with NAD^+^ redox imbalance; however, the hypothesis that NAD^+^ redox imbalance leads to dysfunction of diabetic hearts has not been tested. In this study, we employed mouse models with altered NAD^+^ redox balance to test the hypothesis.

**Methods and Results:** Diabetes was induced in C57BL/6 mice by streptozotocin injections, and diabetic cardiomyopathy (DCM) was allowed to develop for 16 weeks. Diabetic stress led to cardiac dysfunction and lowered NAD^+^/NADH ratio. This diabetogenic regimen was administered to cardiac-specific knockout mice of complex I subunit Ndufs4 (cKO), a model with lowered cardiac NAD^+^/NADH ratio without baseline dysfunction. Cardiac NAD^+^ redox imbalance in cKO hearts exacerbated systolic and diastolic dysfunction of diabetic mice in both sexes. Collagen levels and transcript analyses of fibrosis and extracellular matrix-dependent pathways did not show change in diabetic cKO hearts, suggesting that the exacerbated cardiac dysfunction was likely due to cardiomyocyte dysfunction. We found that cardiac NAD^+^ redox imbalance promoted superoxide dismutase 2 (SOD2) acetylation, protein oxidation, induced troponin I S150 phosphorylation and impaired energetics in diabetic cKO hearts. Importantly, elevation of cardiac NAD^+^ levels by nicotinamide phosphoribosyltransferase (NAMPT) normalized NAD^+^ redox balance, over-expression alleviated cardiac dysfunction and reversed pathogenic mechanisms in diabetic mice.

**Conclusion:** Our results show that NAD^+^ redox imbalance to regulate protein acetylation and phosphorylation is a critical mediator of the progression of DCM, and suggest the therapeutic potential of harnessing NAD^+^ metabolism in DCM.

## Introduction

Diabetes is characterized as a loss of glycemic control, and is a major risk factor of heart failure. Epidemiological studies show a positive correlation of hyperglycemia with the increased risk of heart failure in both Type 1 and Type 2 diabetes patients ^1-3^. Diabetic cardiomyopathy (DCM) is defined as diabetes-associated cardiac dysfunction independent of coronary artery diseases or other confounding cardiovascular diseases ^4^. Mechanisms such as mitochondrial dysfunction and oxidative stress have been identified ^5^. However, how altered metabolism in diabetes, in particular NAD^+^-dependent pathways, leads to cardiac dysfunction is not fully understood.

NAD^+^-dependent pathways have emerged to play critical roles in metabolism-driven disease progression ^6^. NAD^+^ mediates electron transfer and redox reactions in metabolism ^7^. The oxidized form of NAD^+^ (NAD^+^) is converted to the reduced form (NADH) to mediate substrate catabolism (e.g. glycolysis) via many oxidoreductase-dependent metabolic enzymes. The recycling of NADH to NAD^+^, and NAD^+^ redox balance (high NAD^+^/NADH ratio) are essential to maintain substrate catabolism and ATP synthesis. The functions of NAD^+^ have been vastly expanded with the discovery of NAD^+^-consuming enzymes, such as Sirtuins, a family of NAD^+^-dependent deacylases. Sirtuins remove acyl modifications on lysine residues of proteins by turning NAD^+^ into nicotinamide (NAM) and acyl-adenosine diphosphate ribose. Activation of Sirtuins is generally cytoprotective and inhibition of Sirtuins promotes pathogenesis in many metabolic diseases ^8-11^. The availability of NAD^+^ to Sirtuins, via changes in the NAD^+^/NADH ratio or the NAD^+^ pool, is crucial to regulate mitochondrial and cellular functions. To replenish the loss of NAD^+^ consumed by Sirtuins, NAD^+^ is synthesized by several biosynthetic pathways. The NAD^+^ salvage pathway is the dominant pathway used in a mouse heart ^12^: NAM generated from the reactions of Sirtuins is recycled to synthesize nicotinamide mononucleotide (NMN) by nicotinamide phosphoribosyltransferase (NAMPT), the rate-limiting enzyme of the salvage pathway. NMN is then converted into NAD^+^ by nicotinamide mononucleotide adenylyltransferases (NMNATs). Elevating NAD^+^ levels by activation of the NAD^+^ biosynthetic pathways is therapeutic to multiple diseases ^13-20^. We previously showed that NAD^+^-dependent acetylation contributes to pressure overload-induced heart failure, and elevation of NAD^+^ levels alleviates the cardiac dysfunction ^21, 22^. Hyperglycemia is associated with decreased NAD^+^/NADH ratio, and promotes protein hyperacetylation and mitochondrial dysfunction in diabetic organs ^16, 23-27^. Although these data support a hypothesis that NAD^+^ redox imbalance in diabetic hearts promotes cardiac dysfunction, this hypothesis has not been directly tested.

We previously generated a mouse model of mitochondrial dysfunction by deleting Ndufs4, a complex I protein, only in the hearts (cardiac Ndufs4-KO, cKO) ^21^. Complex I deficiency in cKO mice leads to lowered cardiac NAD^+^/NADH ratio, but surprisingly these mice have normal cardiac function and energetics under unstressed condition ^21, 22^. In this study, we challenged this mouse model of latent cardiac NAD^+^ redox imbalance with streptozotocin (STZ)-induced diabetic stress and examined the progression of DCM. We found that cKO mice showed exacerbated cardiac dysfunction in response to diabetic stress. Furthermore, elevation of cardiac NAD^+^ levels with cardiac-specific NAMPT transgenic mice (NAMPT) ^28^ ameliorated the cardiac dysfunction in both diabetic cKO and control mice. The exacerbated cardiac dysfunction in diabetic cKO was associated with increased levels of acetylation of antioxidant enzyme SOD2, oxidative stress and Troponin I (TnI) S150 phosphorylation. Importantly, NAMPT overexpression normalized NAD^+^/NADH ratio in diabetic cKO hearts and reversed these pathogenic mechanisms.

## Research Design and Methods

### Animals care and experiments

All animal care and procedures were approved by the Institutional Animal Care and Use Committee (IACUC) at the Oklahoma Medical Research Foundation and the University of Washington. All procedures were performed in accordance with IACUC regulations. Cardiac-specific Ndufs4 (cKO) mice were generated from the breeding of Ndufs4^flox/flox^ mice (control) with alpha-MHC-Cre expressing mice ^21^. Expression of NAMPT in the heart was achieved by crossing a cardiac-specific NAMPT mouse line with cKO and control mice ^21, 28^. Three to four-month-old littermate mice were used at the beginning of this study. STZ was used to induce beta-cell and insulin depletion, and mice receiving STZ or vehicle were randomly chosen in different cages. STZ in citrate buffer was given to mice intraperitoneally at 50 mg/kg for five days in chronic diabetic experiments. One STZ administration at 50 mg/kg was given in acute STZ experiment. Body weights were measured before, 8 weeks or 15 weeks after STZ injection in different experiments. Blood glucose levels were measured to confirm diabetic phenotypes. STZ-treated mice without hyperglycemia were to be excluded. All diabetic mice reported showed the expected hyperglycemia and no exclusion was needed.

Systolic function and cardiac geometry were assessed by echocardiography with parasternal long axis view and M-mode images were recorded using VEVO 2100 system (VisualSonics) on lightly anesthetized mice. Parameters such as fractional shortening (FS), and left ventricular internal diameter at diastole (LVID;d) were measured, blindly analyzed and calculated by using the software package in the VEVO 2100 system when the heart rate was within 500-600 beats per minute. Using the same system, parameters of diastolic function were measured. E’/A’ ratio was measured by tissue doppler imaging of mitral annulus, e velocity measured by transmitral pulse-wave doppler, and e/E’ ratio was also calculated. Myocardial performance index (MPI) was calculated as the sum of isovolumetric contraction time (IVCT) and isovolumetric relaxation time (IVRT) divided by ejection time (ET). These parameters were measured and calculated by averaging three cardiac cycles.

### Tissue harvest and processing

On the day of tissue harvest, mice were subjected to fasting for 6 hours and anesthetized using pentobarbital. Rib cages were cut open to expose hearts for blood collection by cardiac puncture. Heart weight, wet and dry lung weights, and tibia length were measured post-mortem. Cardiac tissue samples were collected and snap-frozen with liquid nitrogen. Frozen tissues were pulverized using a Tissuelyzer II (Qiagen) for biochemical assays.

### NAD^+^ assay

10-15 mg of pulverized cardiac tissue were used for measurement of NAD^+^ and NADH levels using commercially available kits (BioAssay) under the manufacturer’s instructions.

### Analysis of mRNA levels

Total RNA was extracted from pulverized cardiac tissues using RNeasy fibrous tissue mini kit (Qiagen). RNA concentrations were quantified by nanodrop. To assess expression of genes related to fibrosis, cDNA samples were synthesized, and quantitative PCR reactions were performed using RT2 Profiler PCR Array for mouse extracellular matrix and adhesion molecules genes (Qiagen, Cat. No. 330231 ID: PAMM-013Z) according to manufacturer’s instructions.

### Western blotting

Pulverized cardiac tissues were homogenized in RIPA buffer (Sigma) with protease and phosphatase inhibitor cocktail (Halt, ThermoFisher) and deacetylase inhibitors (10 mM nicotinamide, 10 uM trichostatin A). Protein concentrations of samples were determined by BCA assay (ThermoFisher), and equal amounts of protein (20 ug per sample) were loaded for SDS-PAGE using the Criterion system. Proteins were transferred onto PVDF membrane using Criterion Blotter (Biorad). Blots were blocked in 5% BSA-TBST. Primary antibodies were diluted using 5% BSA-TBST. Antibodies from the following companies were used for Western blot analysis: acetyl-lysine (1:1000, 9441-Cell signaling), SDHA (1:10000, ab14715-Abcam), SOD2 (1:3000, PA5-30604-Thermo Fisher) SOD2-K68Ac (1:10000, ab137037-Abcam), TnI-S150Pi (1:1000, PAS-35410-Thermo), TnI-S23/24Pi (1:1000, 4004S-Cell Signaling), TnI (1:1000, 4002S-Cell Signaling), MyBPC-S282Pi (1:2000, ALX-215-057-R050), MyBPC (1:1000, SC-137237, Santa Cruz), AMPKa (1:1000, 2532S-Cell Signaling) and AMPKa-T172Pi (1:1000, 2535S-Cell Signaling). Protein bands were visualized with chemiluminescence assay (Pierce) with secondary antibodies coupled with HRP using the G-Box imaging system. The protein abundance was analyzed by densitometry with ImageJ.

### Sample analyses by LC-MS/MS

Mouse blood was collected using cardiac puncture at tissue harvest. Blood samples were incubated in EDTA collection tube (BD Diagnostics). After thorough mixing, blood samples were centrifuged at 300 g for five minutes at room temperature. Plasma samples were collected as supernatant and snap-frozen by liquid nitrogen. Glucose levels in plasma were measured by test strips (AlphaTRAK 2 blood glucose tests). Plasma insulin levels were measured by a commercially available kit (Crystalchem).

In metabolite analysis, acetonitrile (ACN), methanol (MeOH), ammonium acetate, and acetic acid, all LC-MS grade, were obtained from Fisher. Ammonium hydroxide was bought from Sigma-Aldrich. PBS was bought from GE Healthcare. The standard compounds were purchased from Sigma-Aldrich and Fisher.

Frozen plasma samples were thawed overnight under 4°C. 500 μL of MeOH and 50 μL of internal standard solution were added to each sample (50 μL) for protein precipitation and metabolite extraction (containing 1,810.5 μM ^13^C_3_-lactate and 142 μM ^13^C_5_-glutamic acid). The mixture was vortexed and stored at -20°C, followed by centrifugation at 14,000 RPM. The supernatants were collected and dried using a CentriVap Concentrator (Labconco). The dried samples were reconstituted with 40% PBS/60% ACN. A pooled sample was used as the quality-control (QC) sample.

The targeted LC-MS/MS method was used in a growing number of studies ^29-31^. Briefly, all LC-MS/MS experiments were performed on an Agilent 1290 UPLC-6490 QQQ-MS system. Each sample was injected twice for analysis using negative and positive ionization mode. Both chromatographic separations were performed in hydrophilic interaction chromatography (HILIC) mode on a Waters XBridge BEH Amide column (Waters). The mobile phase was composed of Solvents A (10 mM ammonium acetate, 10 mM ammonium hydroxide in 95% H_2_O/5% ACN) and B (10 mM ammonium acetate, 10 mM ammonium hydroxide in 95% ACN/5% H_2_O). After the initial 1 min isocratic elution of 90% B, the percentage of Solvent B decreased to 40% at t=11 min. The composition of Solvent B maintained at 40% for 4 min (t=15 min), and then the percentage of B gradually went back to 90%, to prepare for the next injection.

The mass spectrometer is equipped with an electrospray ionization (ESI) source. Targeted data acquisition was performed in multiple-reaction-monitoring (MRM) mode. The extracted MRM peaks were integrated using Agilent MassHunter Quantitative Data Analysis.

### Histology

One to two-millimeter thick heart tissues were cut cross-sectionally at the mid-section and fixed with 4% paraformaldehyde for 24 hours. The tissues were changed to 70% ethanol and fixed tissues were processed for paraffin embedding and sectioning. Tissue sections were stained with trichrome for collagen analysis. The trichrome stained slides were scanned with an Aperio Scanscope AT2 (Leica) at 40x. Quantitative image analysis of whole stained sections was blindly performed using Aperio Brightfield Image Analysis. Toolbox software and a color deconvolution algorithm was used to measure RGB color vectors of blue (positive trichrome stain) and total stained tissue area. The quantitative analysis of cardiac fibrosis for the percentages of collagen (trichrome-blue stain component) and total stained tissue area (mm^2^) was performed. Color images of H&E stained sections were selected at 10X zoom mode using ImageScope and were exported as an image file. Cardiomyocyte sizes were measured using ImageJ ^32^: a masking algorithm was utilized to select cells with clear boundaries. Cross-section fields of cardiomyocytes were chosen and tangential cardiomyocytes were avoided. Further, thresholding of the masked image was used to reduce saturation and set the background level. The threshold was kept similar across all images analyzed. To enhance the cell selection procedure, particle size threshold was set to select cells within a range to exclude bad cells/debris. This helped eliminate cells sectioned tangentially.

## Statistical analysis

For comparisons involving two groups, unpaired 2-tailed t-tests were used. For analysis of sex-dependent changes, two-way ANOVA was used. All analyses were performed using GraphPad Prism 8.0. All data are expressed as mean ± SEM, and a *p*<0.05 was considered significant. Metabolite levels were statistically analyzed by MetaboAnalyst 4.0. Adjusted P-value (FDR) cutoff for the dataset was set at 0.05. Raw data are available as supplementary materials. Heatmaps were generated by Morpheus.

## Results

### 16 weeks of diabetic stress impaired cardiac function and lowered NAD^+^/NADH ratio

Previous studies have shown fragmented evidence for the association of diabetic stress, cardiac dysfunction, and NAD^+^ redox imbalance ^4, 16, 24, 33^. To determine co-existence of cardiac dysfunction and NAD^+^ redox imbalance in diabetic hearts, we induced diabetes using STZ, a drug that promotes beta-cell death and leads to loss of glycemic control. We subjected C57BL/6 wildtype mice (WT) to STZ-induced diabetic stress for 16 weeks, as systolic and diastolic dysfunction has been demonstrated in mice at 16 weeks after STZ treatment ^34^. STZ treatment in WT mice led to significant increase in fasting blood glucose levels at 16 weeks after treatment (Figure 1A). We showed that 16 weeks of diabetic stress in WT mice led to decline in systolic and diastolic function of the heart. Systolic function represented by fractional shortening (FS) decreased in STZ-treated diabetic mice (Figure 1B). E’/A’ ratio decreased, e/E’ ratio and isovolumic relaxation time (IVRT) increased in diabetic mice, indicating diastolic dysfunction (Figure 1C-F). These declines in cardiac function were accompanied by a lowered NAD^+^/NADH ratio (Figure 1G). An *in vitro* study has shown that the effects of STZ are not specific only to the beta cells, and that STZ treatment impairs contractile function of isolated rat cardiomyocytes ^35^. To determine if STZ injection causes acute toxicity in the heart, we examined the acute effects of STZ treatment on the heart. We observed that acute STZ treatment (1 day) did not affect cardiac function, NAD^+^ pool size, and fasting glucose levels (Supp. Figure 1A-D). STZ is rapidly metabolized and excreted by rodents ^36^. Therefore, the lowered NAD^+^/NADH ratio and cardiac dysfunction at 16 weeks after STZ treatment should be caused by chronic diabetic stress (Figure 1A), as observed in other diabetic tissues ^16, 23-27^, but not direct STZ-mediated toxicity to the heart. To determine if the decrease in NAD^+^/NADH ratio precedes cardiac dysfunction, WT mice were subjected to 2-week, STZ-induced diabetic stress. We found that NAD^+^/NADH ratio decreased in hearts after 2-week diabetic stress, while systolic function and diastolic function remained unchanged (Figure 1H-J). The data suggest that NAD^+^ redox imbalance is an early event that precedes cardiac dysfunction in diabetic hearts, and is a potential mechanism that mediates DCM progression.

**Figure 1.**
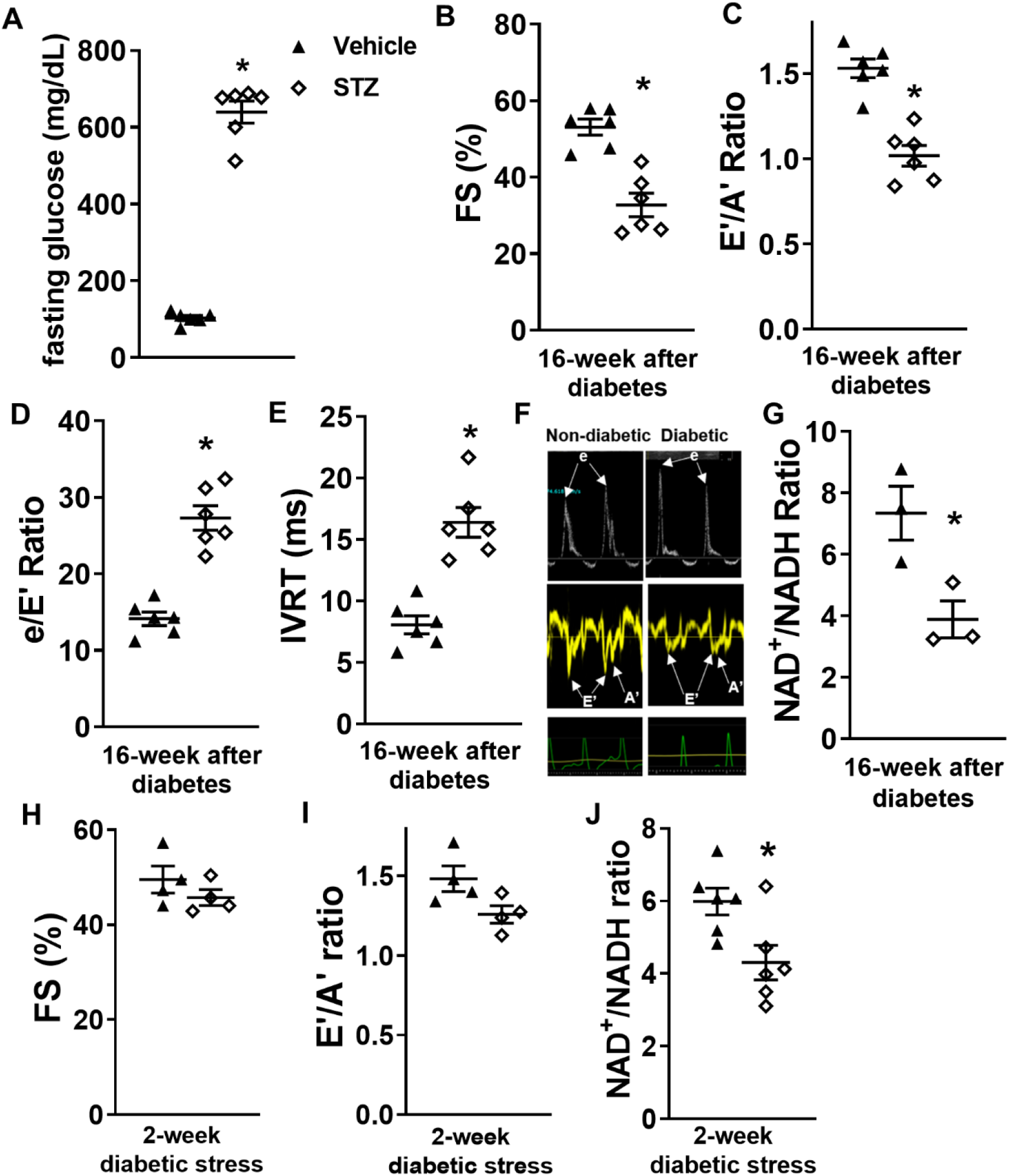
16-week diabetic stress leads to systolic and diastolic dysfunction and lowered NAD^+^/NADH ratio. C57/BL6 wild type (WT) mice were treated with STZ to induce diabetes. Cardiac function was assessed with echocardiography 16 weeks after induction of diabetes. **(A)** Plasma glucose levels were measured 16 weeks after vehicle or STZ treatment. **(B)** Fractional shortening (FS), **(C)** E’/A’, **(D)** e/E’ ratio, and **(E)** isovolumic relaxation time (IVRT) were measured to evaluate systolic and diastolic functions. **(F)** Representative pulsed-wave (upper panels) and tissue (middle panels) doppler images and corresponding electrocardiogram (lower panels) images. N=6. **(G)** Cardiac NAD^+^/NADH ratio were measured. N=3. *: P<0.05 to vehicle. **(H)** FS, **(I)** E’/A’ ratio and **(J)** cardiac NAD^+^/NADH ratio were measured in mice 2-week after diabetic stress. N=4-6. *: P<0.05 to vehicle.

### Latent NAD^+^ redox imbalance exacerbated dysfunction of diabetic hearts

Although diabetic hearts displayed NAD^+^ redox imbalance and dysfunction, the causal role of the lowered NAD^+^/NADH ratio induced by diabetic stress to cardiac dysfunction has not been established. Therefore, we employed cardiac-specific Ndufs4-KO mice (cKO), a mouse model with lowered cardiac NAD^+^/NADH ratio, normal cardiac function and energetics, ^21^ to determine how NAD^+^ redox imbalance changes the progression of DCM. Male control and cKO mice were subjected to 8-week, STZ-induced diabetic stress (Figure 2A). STZ treatment led to insulin depletion, and the extent of insulin depletion was similar in control and cKO mice (Supp. Figure 2A). Fasting blood glucose levels of control and cKO mice at 8 weeks after STZ treatment were up-regulated to similar levels (Supp. Figure 2B). To assess the systemic metabolic changes induced by STZ treatment, we next analyzed plasma samples collected from these mice by quantitative metabolomics. Metabolic stress phenotypes, including increased circulating glucose and ceramide levels, were observed in STZ-induced diabetic control mice, when compared to non-diabetic control mice (Supp. Table 1). Of the 159 aqueous and 85 lipid metabolites surveyed, no significant changes in levels were observed between diabetic control and diabetic cKO mice (Supp. Figure 2C-D, FDR cutoff < 0.05, data available in Supp. Table 2). The results of the plasma metabolomic analyses suggest that cardiac-specific Ndufs4 deletion did not affect peripheral metabolism, and diabetic control and diabetic cKO hearts experienced similar metabolic stress induced by STZ. Despite the similar extent of metabolic stress, diabetic cKO mice exhibited reduced cardiac NAD^+^/NADH ratio compared to diabetic control mice (Figure 2B). The diabetic stress promoted a decline in systolic function of control mice, which was exacerbated in cKO mice 2, 4, and 8 weeks after STZ injection (Figure 2C, Supp. Figure 3A). The diabetic cKO mice also exhibited exacerbated diastolic dysfunction represented by a lowered E’/A’ ratio, increased e/E’ ratio and IVRT, and an elevated myocardial performance index (MPI) compared to diabetic control mice (Figure 2D-G). Left ventricular dilation, cardiac hypertrophy and lung edema did not change in diabetic cKO hearts compared to diabetic controls (Figure 2H-J).

**Figure 2.**
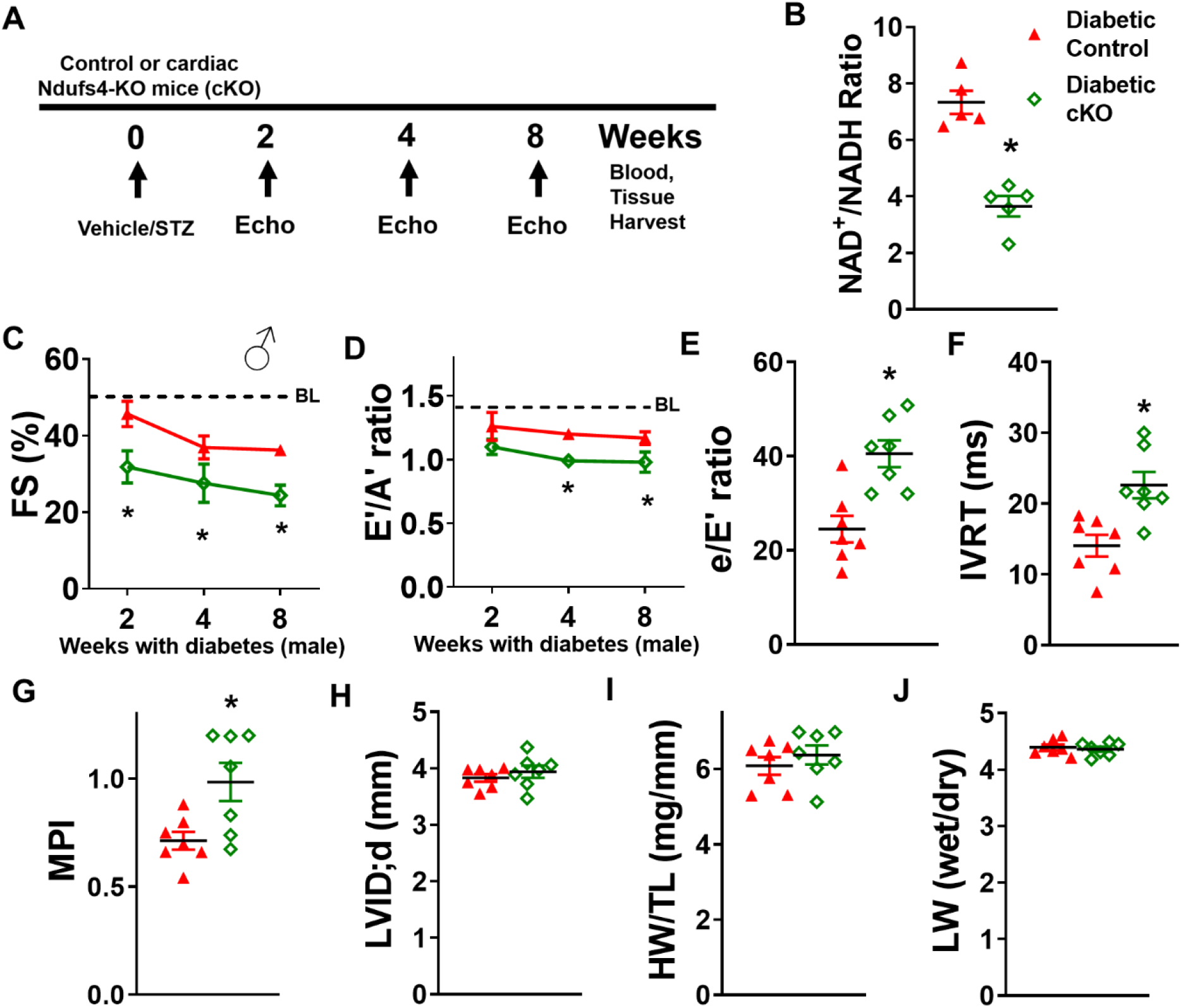
Latent NAD(H) redox imbalance in the heart exacerbates cardiac dysfunctions of diabetic male mice. **(A)** Experimental plan. Control and cKO male mice were treated with STZ to induce diabetes. Cardiac function was measured at 2, 4, and 8 weeks after induction of diabetes. Blood and tissue samples were collected at the endpoint of the study. **(B)** Cardiac NAD^+^/NADH ratio was measured. N=5. Longitudinal changes in **(C)** fractional shortening (FS) and **(D)** E’/A’ ratio of diabetic control or cKO male mice were assessed. **(E)** e/E’ ratio, **(F)** isovolumic relaxation time (IVRT), **(G)** myocardial performance index (MPI) and **(H)** left ventricular internal dimension at diastole (LVID;d) were measured at 8 weeks after diabetes induction. **(I)** Cardiac hypertrophy (heart weight/tibia length; HW/TL) and **(J)** lung edema of diabetic control or cKO mice (lung wet weight/lung dry weight ratio; LW wet/dry) were determined at 8-week endpoint. N=7. *: P<0.05 to diabetic control mice. Dotted lines indicate average baseline (BL) values of non-diabetic control mice.

**Figure 3.**
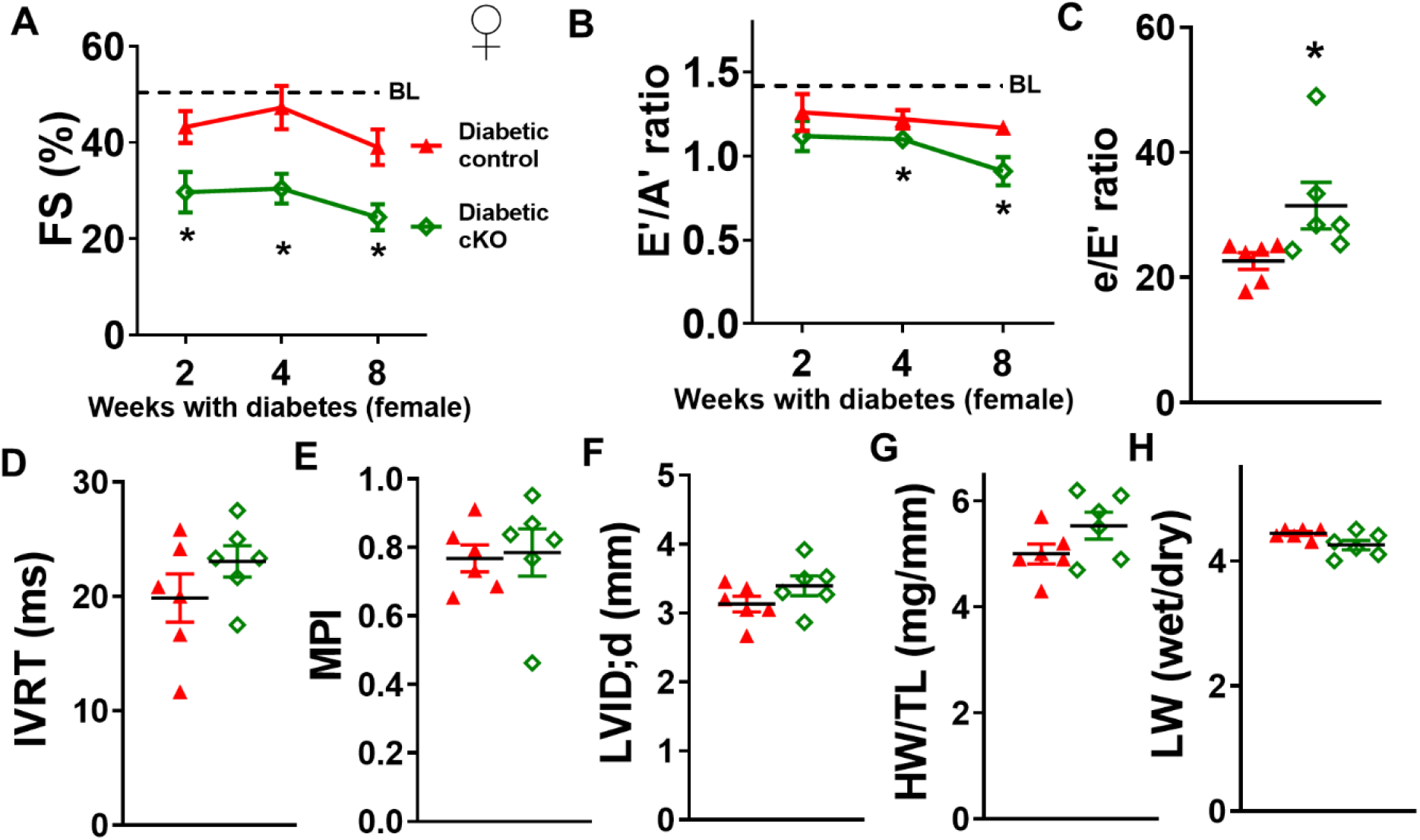
NAD(H) redox imbalance in the heart also exacerbates cardiac dysfunctions of diabetic female mice. Control and cKO female mice were treated with STZ to induce diabetes. Cardiac function was assessed at 2, 4, and 8 weeks after induction of diabetes. **(A)** Longitudinal changes in fractional shortening (FS) and **(B)** E’/A’ ratio of diabetic control or cKO female mice were measured. **(C)** e/E’ ratio, **(D)** IVRT, **(E)** MPI, **(F)** LVID;d, **(G)** HW/TL, and **(H)** LW wet/dry ratio of diabetic control or cKO female mice were determined at 8-week endpoint. N=6. *: P<0.05 to diabetic control mice. Dotted lines indicate average baseline (BL) values of non-diabetic control mice.

We performed the same diabetogenic protocol (Figure 2A) on female control and cKO mice. Similarly, diabetic stress induced systolic and diastolic dysfunction in female diabetic control mice, which was significantly worsened in female diabetic cKO mice (Figure 3A-C, Supp Figure 3B). IVRT, MPI, left ventricular dilation, cardiac hypertrophy, and lung edema were not different between female diabetic control and cKO mice (Figure 3D-H). To determine if cardiac-specific Ndufs4 deletion mediates any sex-dependent changes on DCM progression, we compared the effects of STZ on cardiac function and geometry of male and female diabetic control and cKO mice. Although Ndufs4 deletion only significantly worsened IVRT and MPI in male diabetic cKO mice but in not female diabetic cKO mice, there were no significant interactions between sex and genotype by two-way ANOVA (Supp. Table 3). Overall, the results suggest that NAD^+^ redox imbalance is a positive regulator for the progression of DCM in both sexes.

### Tissue fibrosis was not associated with exacerbated DCM induced by NAD^+^ redox imbalance

Diabetic hearts are often associated with increased fibrosis, which contributes to cardiac dysfunction, especially diastolic dysfunction ^37^. We therefore quantified tissue collagen contents of mid-sectioned heart slices by trichrome staining. We observed similar fibrosis levels (∼4%) in diabetic control and diabetic cKO hearts (Figure 4A). There was also no difference in cardiomyocyte size between the two groups (Figure 4B). Consistent with the fibrosis levels, expression of extracellular matrix (ECM)-related genes were similar between diabetic control and diabetic cKO hearts: transcript levels of Adamts proteinases, integrins, laminins, matrix metalloproteinases and collagen subtypes (Figure 4C-D, Supp. Figure 4A-C) were not different. These results suggest that altered ECM environment and fibrosis do not account for the exacerbated dysfunction, but cardiomyocyte dysfunction may be the culprit.

**Figure 4.**
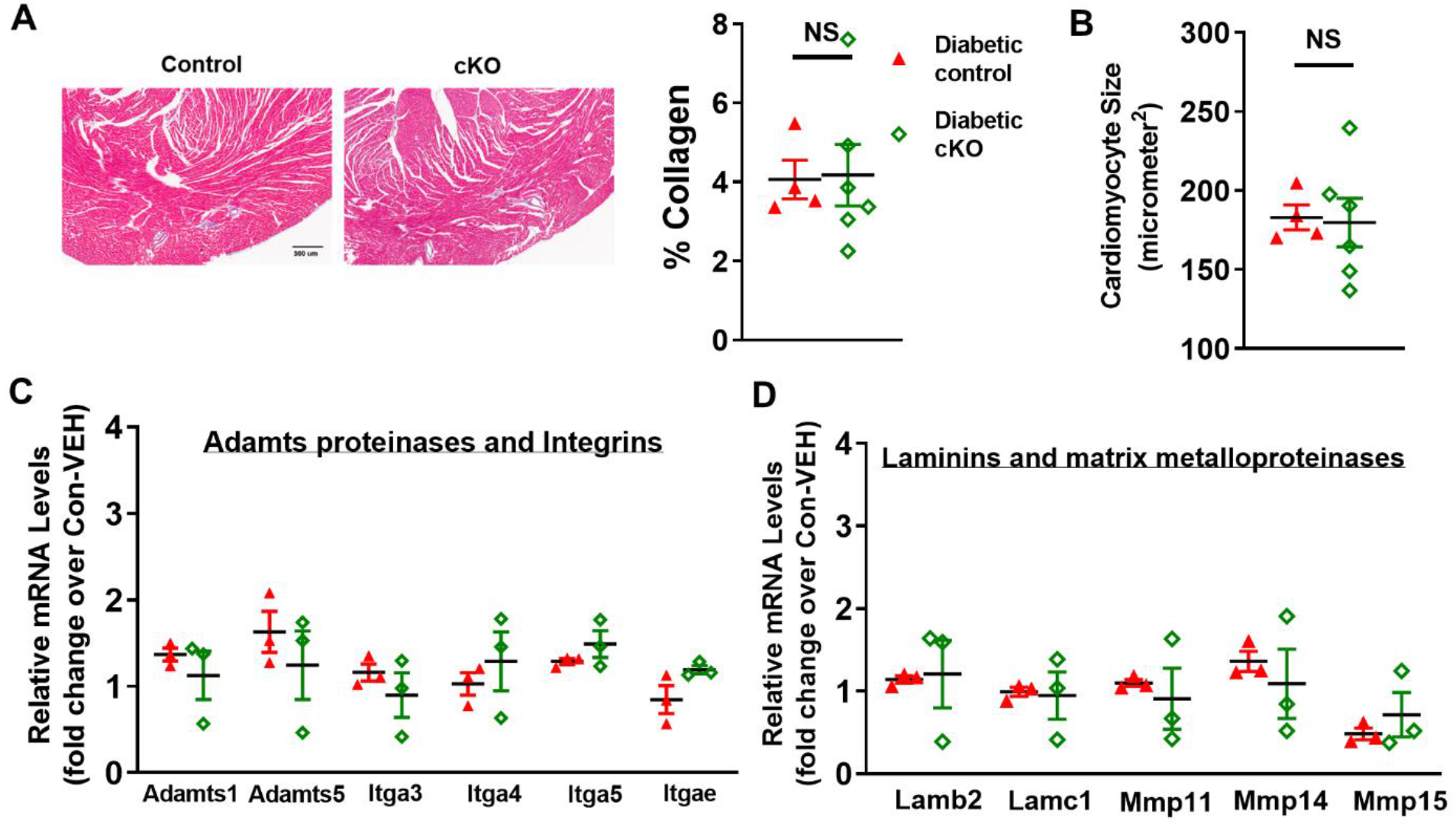
NAD^+^ redox imbalance exacerbates diabetic cardiomyopathy, independent on tissue fibrosis. **(A)** Collagen levels of diabetic mouse hearts were quantified by trichrome staining. **(B)** Cardiomyocyte sizes of these hearts were quantified. N=4-6. Transcript expression levels of fibrosis-related genes were quantified by qPCR analyses. **(C)** Adamts proteinases and integrins, and **(D)** laminins and MMPs in diabetic control and diabetic cKO male hearts were measured. N=3. *: P<0.05 to diabetic control mice.

### Elevated SOD2 acetylation promoted oxidative stress in diabetic cKO hearts

NAD^+^ is required for deacetylation by Sirtuins, and NAD^+^ redox imbalance has been associated with protein hyperacetylation. Therefore, we explored the roles of NAD^+^ redox imbalance in regulating protein acetylation in diabetic hearts. Consistent with the lowered NAD^+^/NADH ratio, global protein acetylation increased in diabetic cKO hearts compared to diabetic controls (Figure 5A, Supp. Figure 4D). Specifically, we observed that acetylation levels of superoxide dismutase 2 at lysine-68 (SOD2-K68Ac) were elevated in diabetic cKO hearts (Figure 5B, Supp. Figure 4E). SOD2-K68Ac is known to suppress its antioxidant activity ^38^, suggesting that diabetic cKO hearts could be more prone to oxidative stress. Consistently, levels of protein carbonylation, an irreversible protein oxidative modification used as a marker of oxidative stress, were elevated in diabetic cKO hearts (Figure 5C, Supp. Figure 4F). Transcript levels of NADPH oxidase (Nox) isoforms, key ROS generating proteins, were similar in diabetic control and diabetic cKO hearts (Figure 5D). These results suggest that NAD^+^ redox imbalance promotes oxidative stress through SOD2 acetylation in diabetic hearts.

**Figure 5.**
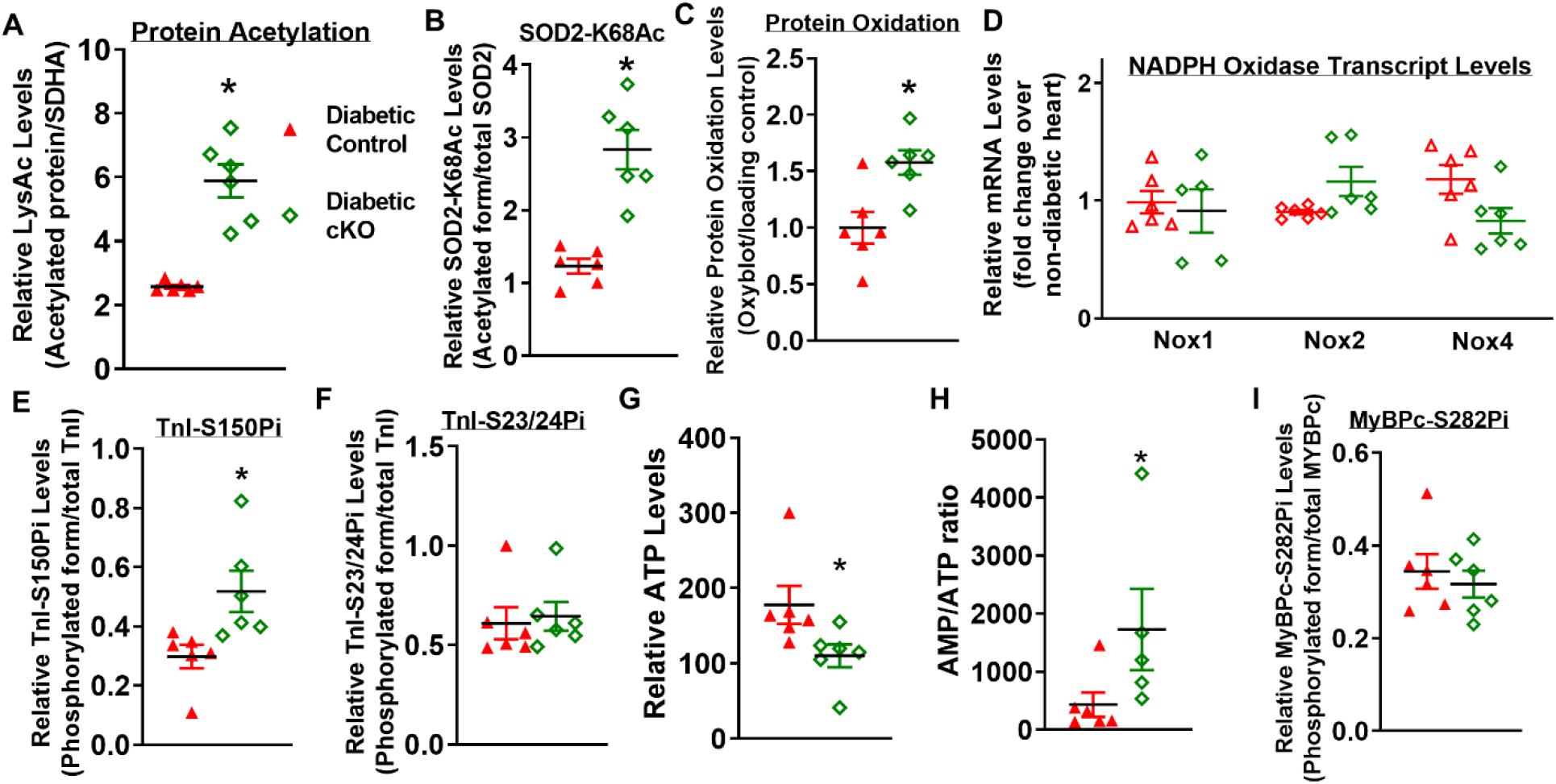
NAD^+^ redox imbalance regulated SOD2 acetylation, protein oxidation and myofilament protein phosphorylation. **(A)** Global lysine acetylation levels of protein extracts from diabetic control or diabetic cKO hearts were assessed by Western blot. **(B)** Acetylation levels of SOD2 at lysine-68 were assessed by Western blot analysis. **(C)** Protein oxidation levels of diabetic mouse hearts were determined by Oxyblot analysis. **(D)** Relative mRNA levels of pro-oxidant genes (Nox1, Nox2 and Nox4) were measured by qPCR. Phosphorylation levels of **(E)** TnI at Serine 150 (TnI-S150Pi), **(F)** TnI-S23/24Pi, **(G)** ATP levels and **(H)** AMP/ATP ratio were measured. Phosphorylation levels of **(I)** MyBPc at Serine 282 (MyBPc-S282Pi) were measured. N=6. *: P<0.05 to diabetic control mice.

### NAD^+^ redox imbalance regulated TnI phosphorylation

We then examined if phosphorylation of myofilament proteins, which control cardiomyocyte contraction and relaxation, were altered by NAD^+^ redox imbalance in diabetic hearts. Phosphorylation states of troponin I (TnI) and myosin binding protein c (MyBPc) can modulate contractile and relaxation properties of the myocardium in diseased hearts ^39, 40^. TnI is an inhibitory subunit of troponin, and phosphorylation of TnI at S150 and at S23/24 (TnI-S150Pi and TnI-S23/24Pi) has been shown to coordinately regulate myofilament calcium sensitivity as an adaptive response in ischemic hearts ^41, 42^. Diabetic cKO hearts displayed increased levels of TnI-S150Pi compared to diabetic controls, while levels of TnI-S23/24Pi were not different (Figure 5E-F, Supp Figure 4G-H). As TnI-S150Pi prolongs calcium dissociation ^41^, these results suggest that NAD^+^ redox imbalance induces TnI-S150 phosphorylation to exacerbate diastolic dysfunction in DCM (Figure 2D-F). AMPK phosphorylates TnI-S150, and is activated by lowered cellular energy states ^43^. Despite normal energetics in unstressed cKO hearts ^21^, we found that ATP levels decreased and AMP/ATP ratio increased in diabetic cKO hearts, compared to diabetic controls (Figure 5G,H). AMP and ADP levels trended to increase in diabetic cKO hearts (Supp. Figure 4I,J). These results indicated lowered cellular energy status in diabetic cKO hearts compared to diabetic controls. AMPK-T172 phosphorylation is also known to activate AMPK, but was not different between diabetic control and diabetic cKO hearts (Supp. Figure 4K). The results suggest that impaired energetics in diabetic cKO hearts mediates AMPK activation and the increased phosphorylation of TnI-S150 for the exacerbated dysfunction.

MyBPc is a sarcomeric protein that interacts with myosin and actin, and phosphorylation of MyBPc fine-tunes actin-myosin cross-bridge cycling to regulate cardiac contraction and relaxation. MyBPc S282 phosphorylation (MyBPc-S282Pi) is critical for normal cardiac function, and reduced MyBPc-S282Pi has been observed in failing hearts ^40^. Levels of MyBPc-S282Pi were unchanged in diabetic cKO hearts (Figure 5I, Supp. Figure 4L), suggesting that it is not responsible for the worsened cardiac function.

### Elevation of NAD^+^ levels ameliorated dysfunction of diabetic hearts

To validate the causal role of cardiac NAD^+^ redox imbalance in DCM, we examined the effects of elevation of cardiac NAD^+^ levels by cardiac NAMPT over-expression on DCM development. Transgenic mice expressing NAMPT in the heart (NAMPT) were crossed with cKO or control mice ^22^. The same experimental protocol was used on these mice (Figure 6A). Over-expression of NAMPT in the heart increased NAD^+^/NADH ratio and NAD^+^ pool in diabetic cKO hearts, and improved systolic and diastolic dysfunction (Figure 6B-D, Supp Figure 5A). These improvements occurred despite the similar extents of hyperglycemia in cKO and cKO:NAMPT mice (Supp. Figure 5B). The body weight changes at 8 weeks after diabetes induction were similar (Supp. Figure 5C). Heart rate, left ventricular dilation, hypertrophy, and lung edema were also unchanged, while IVRT and MPI were improved by cardiac-specific NAMPT over-expression (Supp. Figure 5D-I). Importantly, NAMPT over-expression lowered the levels of SOD2-K68Ac and TnI-S150Pi in diabetic cKO hearts, while TnI-S23/24Pi levels were similar (Figure 6E-G, Supp. Figure 5J-L). To determine if preventing NAD^+^ redox imbalance also ameliorates DCM progression in control mice, we compared the impacts of diabetic stress on cardiac function of cardiac-specific NAMPT transgenic mice and their control littermates. We showed that diabetic NAMPT mice have higher fractional shortening and E’/A’ ratio compared to diabetic control mice, suggesting ameliorated systolic and diastolic dysfunction by NAMPT over-expression (Figure 6H,I). These results collectively support the causal roles of NAD^+^ redox imbalance in the progression of DCM.

**Figure 6.**
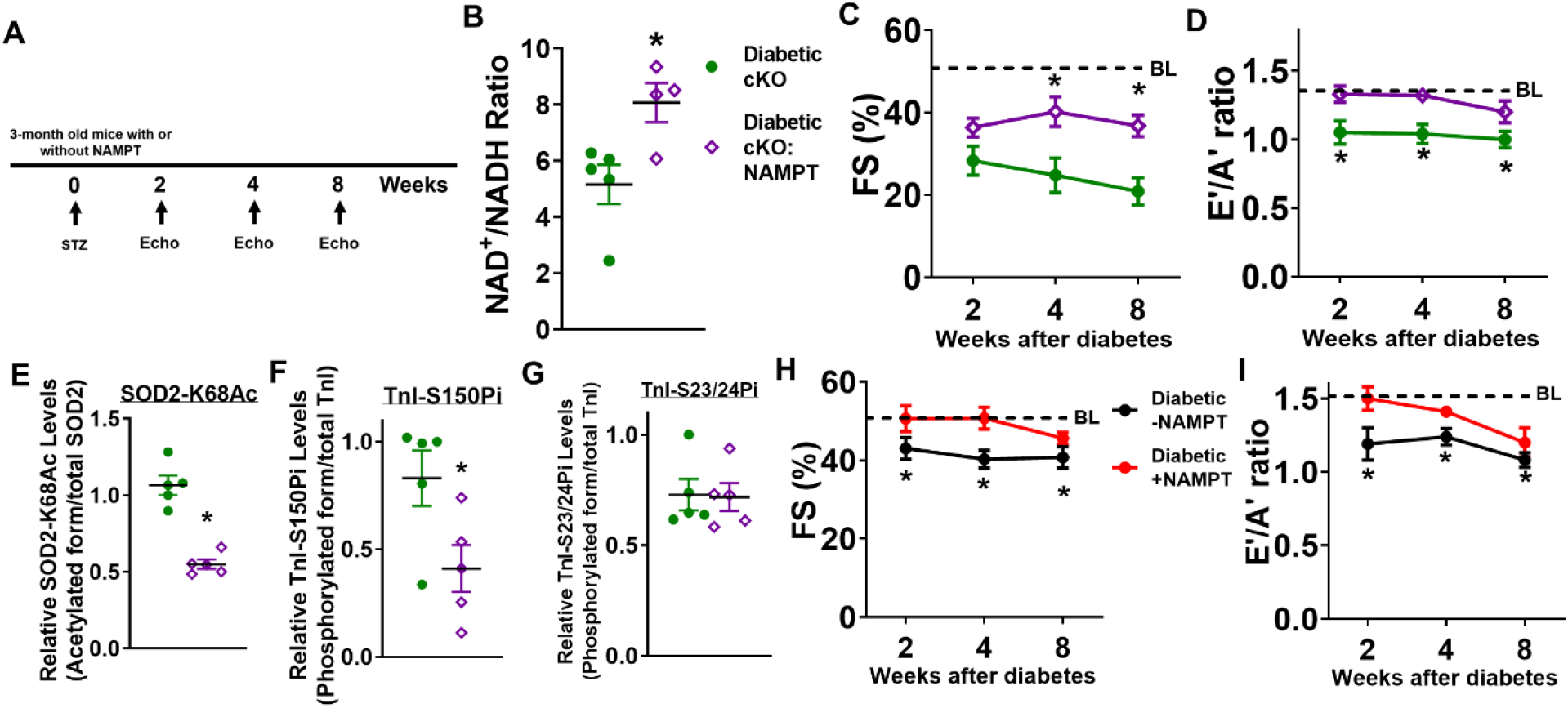
Elevation of cardiac NAD^+^ levels alleviates diabetic cardiomyopathy in cKO and control mice. **(A)** Experimental plan. STZ was administered to male cKO and cKO:NAMPT mice to induce diabetes. Cardiac function of cKO and cKO:NAMPT mice was measured at 2, 4, and 8 weeks after induction of diabetes. **(B)** Cardiac NAD^+^/NADH ratio were measured. N=4-5. **(C)** FS and **(D)** E’/A’ ratio were assessed longitudinally. N=6. Levels of **(E)** SOD2-K68Ac, **(F)** TnI-S150Pi, and **(G)** TnI-S23/24Pi of indicated hearts were measured by Western blots. N=5. **(H)** FS and **(I)** E’/A’ ratio were measured longitudinally in control mice with or without NAMPT expression in the hearts after diabetes induction. N=5. *: P<0.05 to diabetic cKO mice or diabetic-NAMPT mice.

## Discussion

Diabetes increases the risk of heart failure and causes DCM. Although pathogenic mechanisms such as lipotoxicity, oxidative stress and mitochondrial dysfunction have been demonstrated in diabetic hearts ^5^, the mechanisms by which diabetes-induced metabolic stresses lead to cardiac dysfunction are not fully understood. Hyperglycemia and dyslipidemia are the hallmarks of diabetes, and diabetes alters cardiac metabolism such as substrate utilization ^5^. NAD^+^ is an important redox cofactor for glucose and lipid oxidation. Changes in NAD^+^ metabolism such as NAD^+^ redox imbalance emerge as critical mediators of disease pathogeneses ^21, 22^. NAD^+^ redox imbalance has long been observed in tissues from diabetic mice ^16, 23-25^, but the hypothesis that diabetes-driven NAD^+^ redox imbalance promotes DCM remains to be tested. In this study, we demonstrated the causal role of NAD^+^ redox imbalance in promoting DCM. The key findings in supporting this notion are that 1) cardiac NAD^+^/NADH ratio decreases and precedes cardiac dysfunction after diabetic stress; 2) a pre-existing, lowered cardiac NAD^+^/NADH ratio exacerbates the progression of DCM, which is alleviated by elevating NAD^+^ levels; and 3) NAD^+^ redox imbalance mediates dysfunction of diabetic hearts in part by increased oxidative stress and TnI-S150 phosphorylation, but not by enhanced cardiac fibrosis.

Diabetic stress is associated with lowered NAD^+^/NADH ratio, also known as pseudohypoxia, in various tissues including hearts ^24, 44^. Functional decline in diabetic hearts has also been reported in parallel ^5^. However, whether NAD^+^ redox imbalance causes DCM has not been directly tested. We found that 16-week diabetic stress lowered cardiac NAD^+^/NADH ratio, accompanied by systolic and diastolic dysfunction. We also showed that the decline in cardiac NAD^+^/NADH ratio occurred before cardiac dysfunction. Importantly, we observed that a mouse model with lowered cardiac NAD^+^/NADH ratio (due to Ndufs4 deficiency in hearts) exhibits exacerbated DCM, and that cardiac dysfunction in diabetic cKO or control hearts can both be rescued by restoration of NAD^+^/NADH ratio (Figure 2,3,6). This is the first direct evidence to support the causal role of NAD^+^ redox imbalance in the progression of DCM. A recent report showed the role of hepatic NAD^+^ reductive stress (low NAD^+^/NADH ratio) in regulating changes in diabetes-induced metabolite levels (alpha-hydroxybutyrate) and insulin resistance in vivo ^45^, supporting the pathogenic role of NAD^+^ redox imbalance in diabetes and its complications. We previously demonstrated that NAD^+^ redox imbalance is a critical mediator in pressure overload-induced heart failure ^22^. Whether diabetes and hypertension have additive or synergistic effects on cardiac NAD^+^ redox imbalance to promote cardiac dysfunction remains to be determined.

While the causal role of NAD^+^ redox imbalance for DCM is well-supported by our current data, the mechanism by which diabetic stress induces NAD^+^ redox imbalance is not well-established. Previous studies have provided evidence that hyperglycemia and dyslipidemia promote NAD^+^ redox imbalance ^27, 46^. Mitochondrial dysfunction and impaired NAD^+^-dependent malate aspartate shuttle (MAS) have been shown as important mediators of the lowered NAD^+^/NADH ratio ^27, 47, 48^. MAS delivers cytosolic NADH to electron transport chain for ATP generation. Interestingly, MAS activity is suppressed in hypertrophied hearts in a NAD^+^ redox balance-dependent manner ^22, 49, 50^. Whether MAS activity decreases in diabetic hearts and leads to NAD^+^ redox imbalance requires further investigations.

Fibrosis is a hallmark of heart disease of different etiologies, especially in diabetic hearts ^37^, and we did observe a slight increase in cardiac fibrosis in STZ-treated diabetic hearts (Supp. Figure 4M). However, we did not observe increased fibrosis associated with exacerbated dysfunction of diabetic cKO hearts. The similar extents of fibrosis in diabetic control and diabetic cKO hearts suggest that other mechanisms, e.g. cardiomyocyte dysfunction, is responsible for the exacerbated DCM by NAD^+^ redox imbalance. Our results suggest that the exacerbated DCM progression can be mediated by cardiomyocyte dysfunction induced by elevated SOD2 acetylation and oxidative stress in diabetic cKO hearts. Other NAD^+^-sensitive protein acetylation, e.g. sensitized permeability transition pore by acetylation, may also play a role in DCM progression ^22^.

Our data showed that diabetic cKO hearts displayed elevated TnI-S150Pi, exacerbated diastolic dysfunction and prolonged IVRT, which are all restored by NAMPT over-expression. AMPK-dependent TnI-S150Pi increases myofilament calcium sensitivity and prolong calcium dissociation ^39, 41, 42^, while TnI-S23/24Pi decreases calcium sensitivity and accelerates calcium dissociation ^39^. Combination of TnI-S150Pi and TnI-S23/24Pi retains calcium sensitivity and accelerates calcium dissociation, playing an adaptive role during ischemic injury ^39^. Increased TnI-S150Pi and unchanged TnI-S23/24Pi in diabetic cKO hearts would suggest that NAD^+^ redox imbalance promotes TnI-S150Pi to play a maladaptive role in diastolic dysfunction. Although unstressed cKO hearts have normal energetics ^21^, we detected impaired energetics in diabetic cKO hearts (Figure 5G,H). The reduced cellular energy status can activate AMPK and in turn augment TnI-S150Pi. These results suggest that NAD^+^ redox imbalance predisposes cKO hearts to diabetic stress by promoting nucleotide-dependent AMPK activation and TnI-S150Pi.

We showed that cardiac NAMPT over-expression improved cardiac function in diabetic cKO mice, supporting that NAD^+^ redox imbalance is responsible for the exacerbated DCM progression. The observation that cardiac NAMPT over-expression also improved cardiac function in diabetic control mice without Ndufs4 deletion further supports a causal role of NAD^+^ redox imbalance in DCM progression. NAMPT over-expression in cardiomyocytes has been shown to promote hypertrophy and inflammation via secreted form of NAMPT (eNAMPT) ^51^. In this study, we did not observe significant differences in eNAMPT levels in plasma of mice with or without NAMPT over-expression (Supp. Figure 5M). Therefore, eNAMPT and its role in cardiac hypertrophy and inflammation should not play a role in the interpretation of this study. It is conceivable that disease mechanisms of DCM involve other NAD^+^-dependent pathways beyond NAD^+^ redox imbalance and protein acetylation reported here. NAD^+^ metabolism involves metabolites and enzymes in the NAD^+^ consumption, synthesis and redox pathways, coordinating cellular NAD^+^ homeostasis ^52^. Our study employed cardiac-specific perturbations of NAD^+^ redox balance (cKO and NAMPT) and demonstrated the roles of cardiac NAD^+^ redox imbalance in DCM progression. However, we cannot rule out the contributions of NAD^+^-derived metabolites and systemic changes in NAD^+^ metabolism (e.g. plasma NAD^+^ metabolites) in promoting DCM progression. Metabolites in the NAD^+^ salvage pathway, e.g. nicotinamide mononucleotide or nicotinamide riboside, are prime agents for pharmacologic elevation of NAD^+^ levels as therapeutics for heart disease ^22, 53^. In addition, the benefit of activation of NAD^+^ synthesis by harnessing the de novo pathway and the role of NMRK2 up-regulation in cardiomyopathy have been reported ^53, 54^. Therefore, further assessments of changes in NAD^+^ metabolism in DCM are warranted, and will likely identify new targets for therapy. A limitation of this study is that we employed a STZ-induced Type 1 diabetic model, and the roles of NAD^+^ redox imbalance in Type 1 DCM progression cannot be generalized to DCM in Type 2 diabetes models (e.g. high-fat diet feeding) or cardiac dysfunction caused by other cardiac risk factors such as aging. High-fat diet feeding and aging have been associated with abnormal NAD^+^ metabolism ^16, 55^; however, specific mechanisms that alter NAD^+^ metabolism (redox state, synthesis or consumption) in these conditions and their roles in cardiac dysfunction require further investigations.

This study establishes the causal role of NAD^+^ redox imbalance in DCM. Our results support that cardiac-specific NAD^+^ redox imbalance mediates cardiac dysfunction in DCM in both sexes. This highlights the importance to identify adaptive and maladaptive mechanisms triggered by NAD^+^ redox imbalance and altered NAD^+^ metabolism in DCM. Our pre-clinical data also demonstrate the potential benefit of expanding the NAD^+^ pool as a therapy for DCM.

## Acknowledgements

CFL designed and performed the experiments, interpreted the data, and wrote the manuscript. YAC performed experiments and wrote the manuscript. ADC, XS, HG, and CML performed the experiments and analyzed the data. CFL is the guarantor of the study and has full access to all the data in the study and takes responsibility for the integrity of the data and accuracy of the data analysis.

RT and JS kindly provided mouse models. All authors have reviewed and edited the manuscript.

---

**Supplementary Figure 1.**
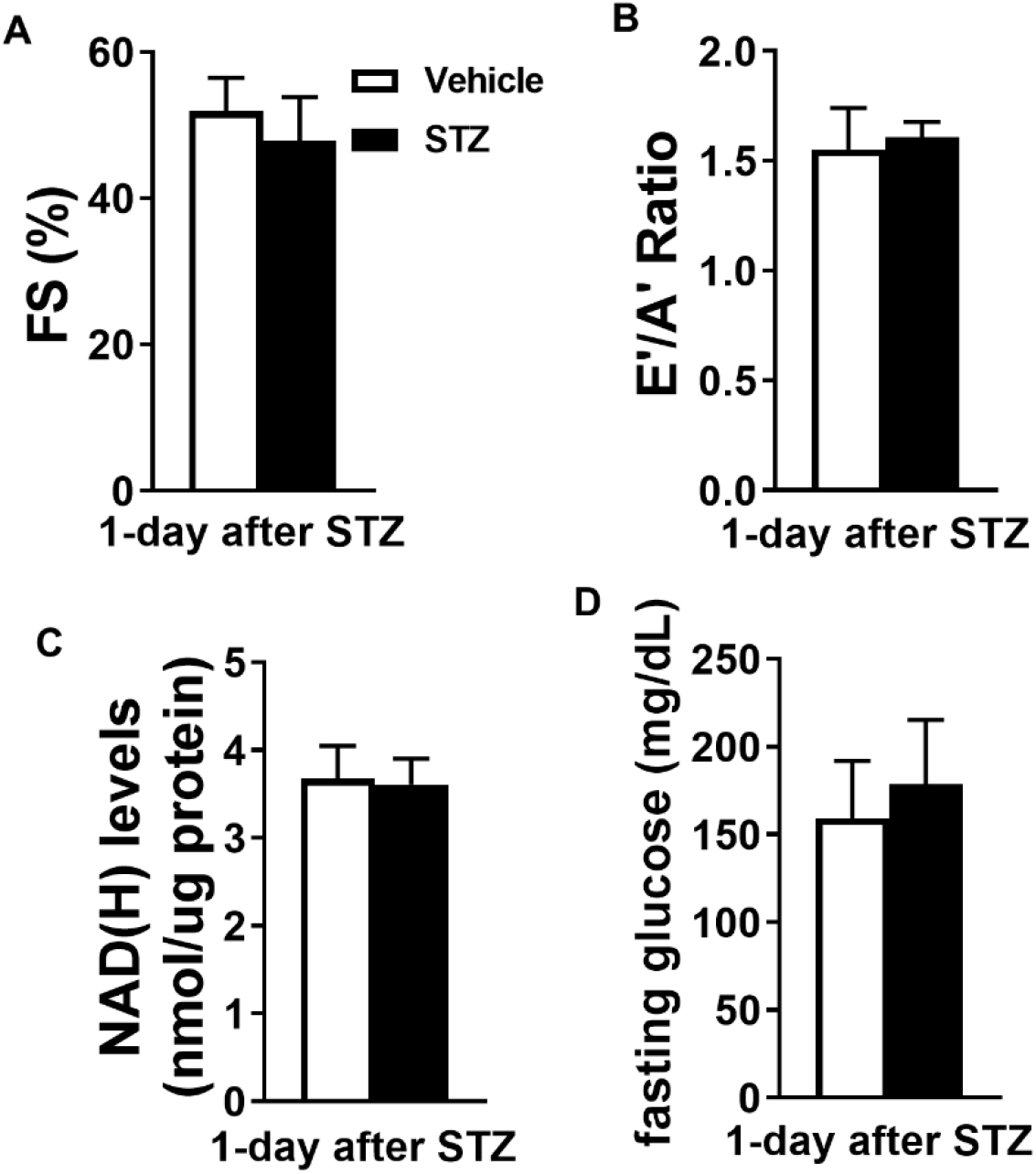
**(A)** FS, **(B)** E’/A’ ratio and **(C)** NAD(H) pool and **(D)** fasting glucose levels were measured in mice 1-day after STZ injection.

**Supplementary Figure 2.**
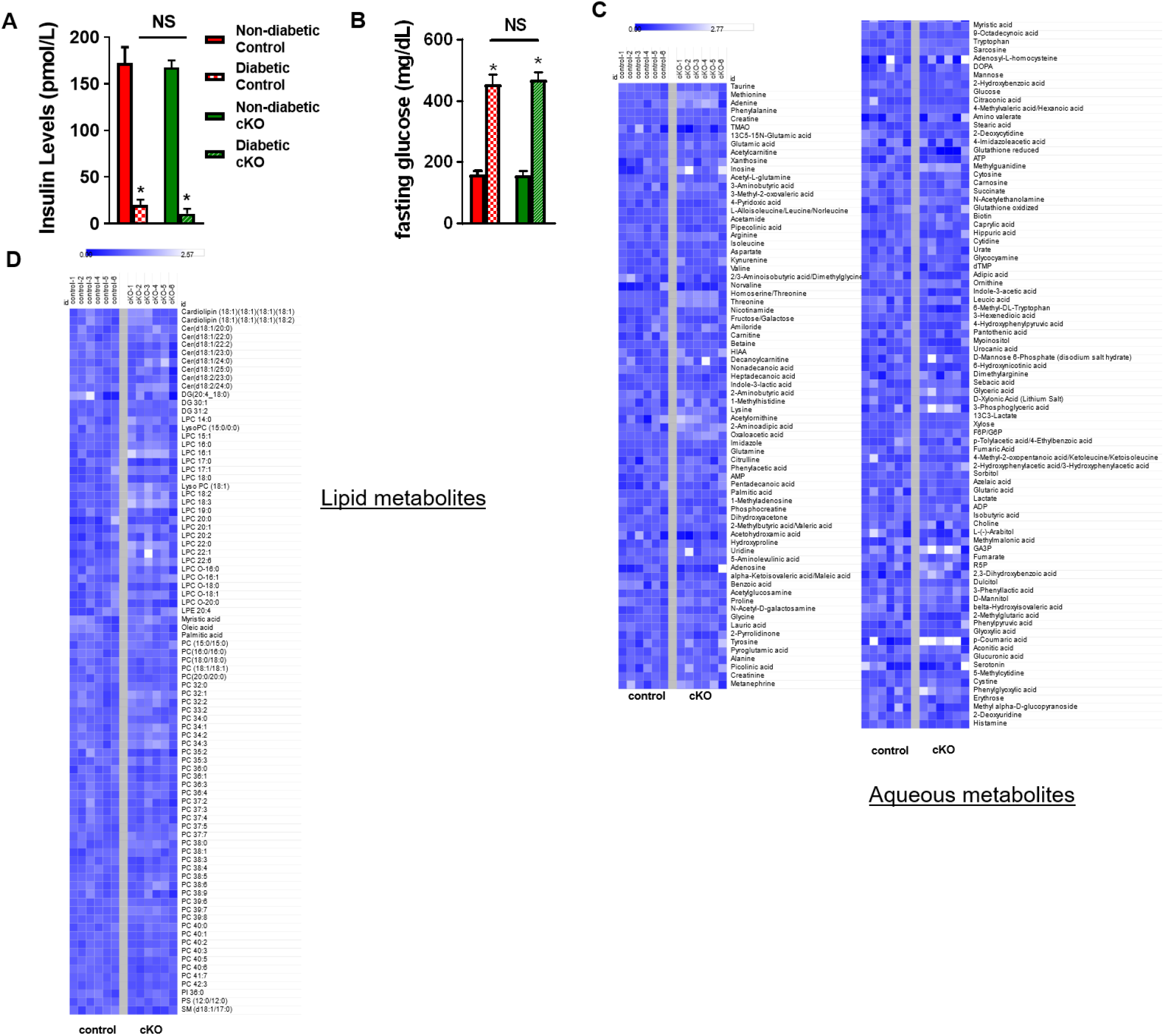
**(A)** Plasma insulin levels and **(B)** fasting glucose levels of male mice with indicated treatment were measured. Heatmaps of **(C)** aqueous and **(D)** lipid metabolite levels in plasma of diabetic control and diabetic cKO mice. Heatmaps were generated using Morpheus. Raw data can be found in supplementary Table 2. N=6 male mice. *: P<0.05 to corresponding non-diabetic group.

**Supplementary Figure 3.**
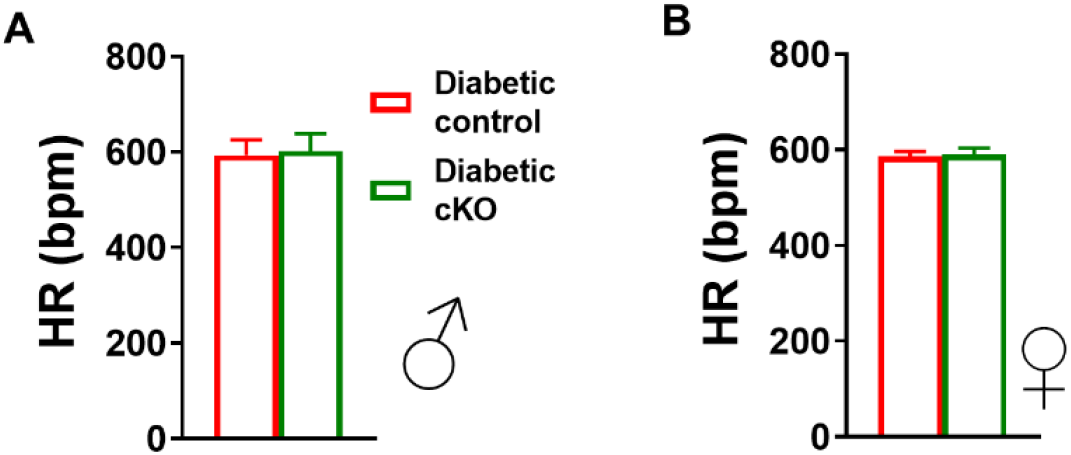
Heart rates (HR) of **(A)** male, **(B)** female cohorts of diabetic control and diabetic cKO mice. N=6-7.

**Supplementary Figure 4.**
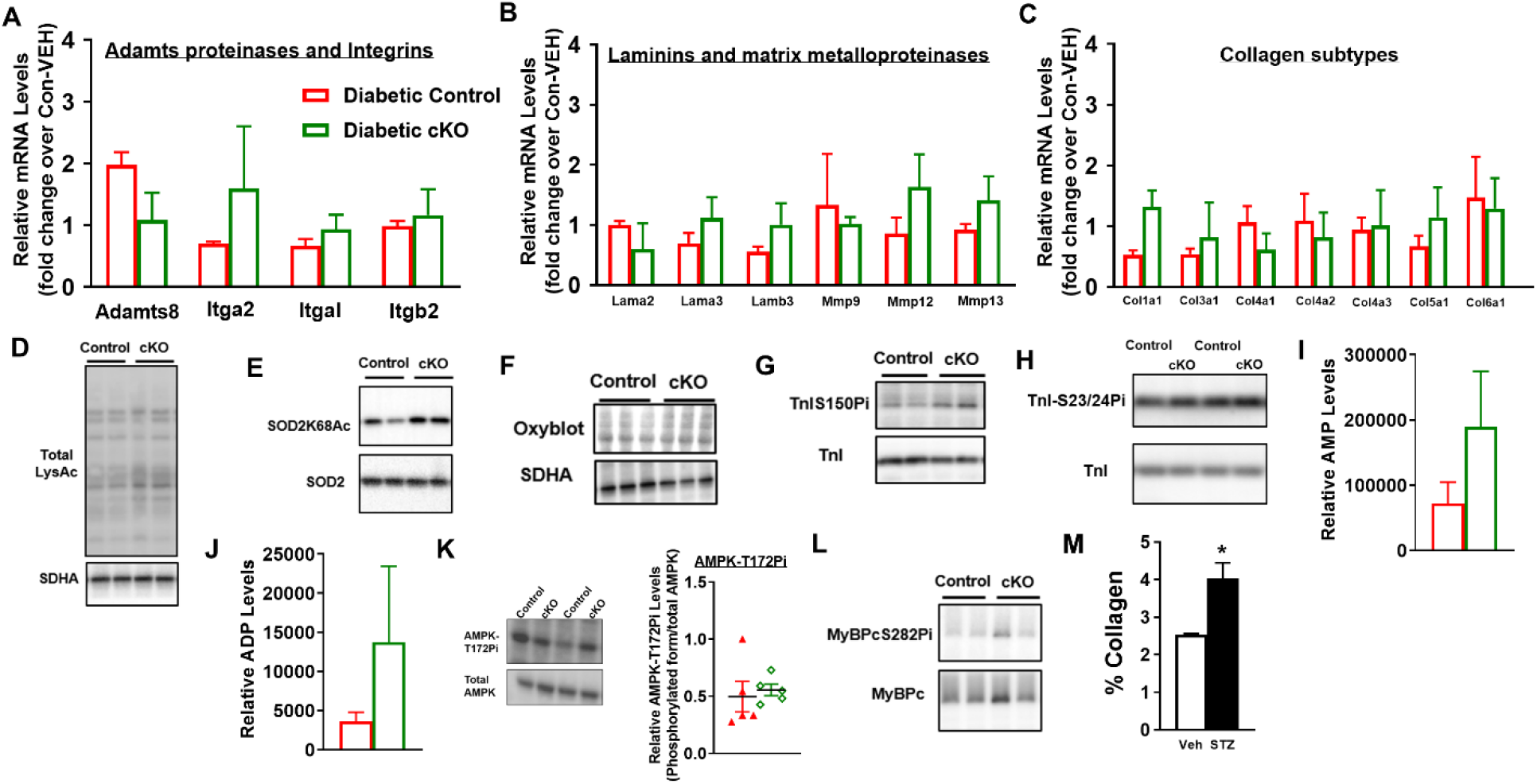
Additional gene expression analysis of **(A)** Adamts proteinases and integrins, **(B)** laminins and MMPs, and **(C)** collagen subtypes in diabetic control and diabetic cKO hearts were performed. N=3 male mice. **(D-H)** Representative Western blots for results shown in Figure 5. **(I)** AMP and **(J)** ADP levels were quantified in control diabetic and cKO diabetic hearts. Phosphorylation levels of **(K)** AMPK-T172 and **(L)** MyBPc-S282 were measured. **(M)** Collagen levels in non-diabetic and STZ-induced diabetic hearts.

**Supplementary Figure 5.**
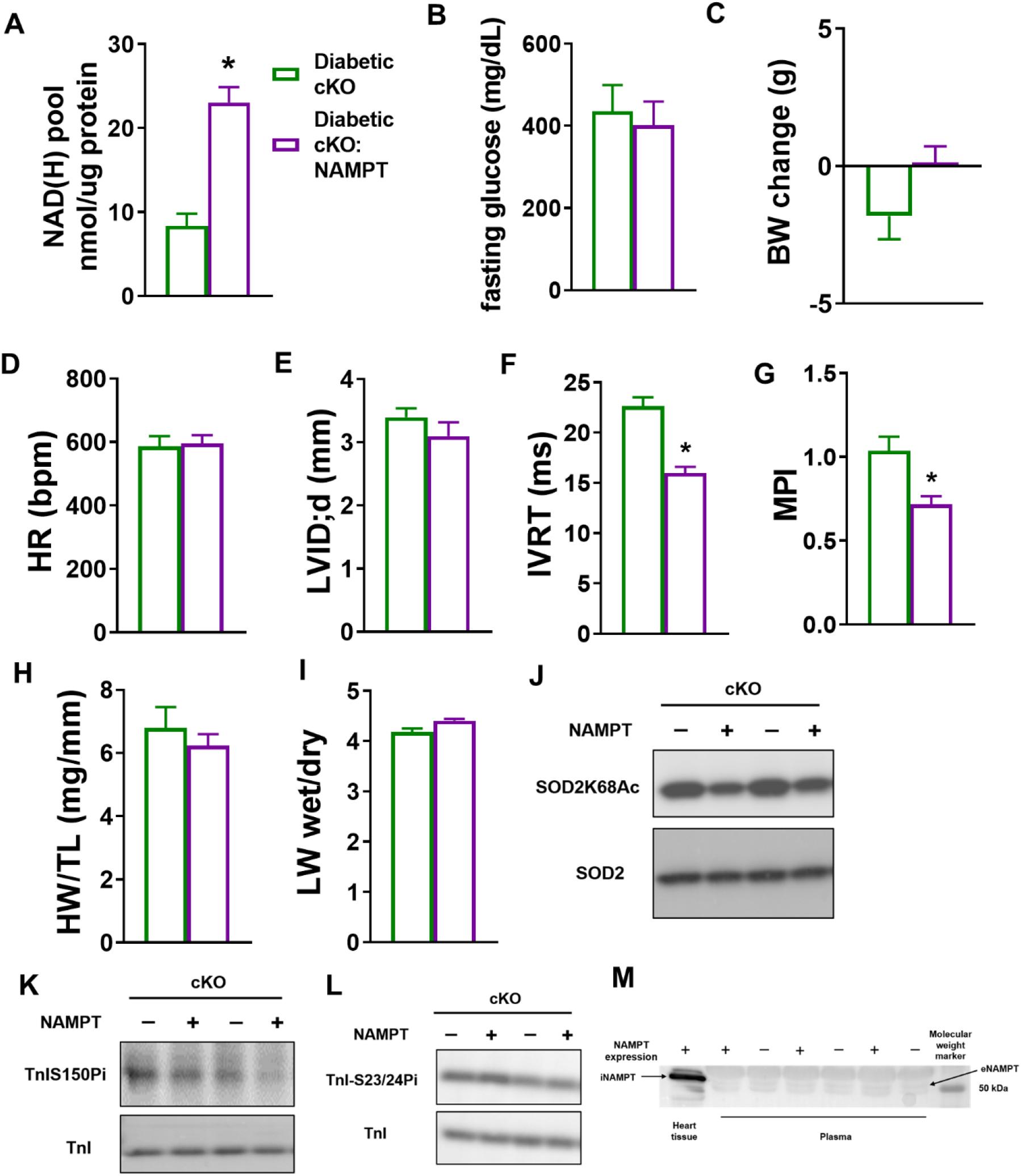
**(A)** NAD^+^ pool of diabetic cKO and diabetic cKO:NAMPT mice 8 weeks after STZ treatment were measured. N=5 male mice. **(B)** Fasting glucose levels, **(C)** body weight (BW) change, **(D)** heart rate (HR) at the time acquiring echocardiography data, **(E)** LV dilation, **(F)** IVRT, **(G)** MPI, **(H)** hypertrophy and **(I)** lung edema of indicated mice were measured at 8-week endpoint. **(J-L)** Representative Western blots for results shown in Figure 6. **(M)** Levels of plasma NAMPT protein were measured by Western blots. *: P<0.05 to diabetic cKO mice.

